# The combination of three CD4-induced antibodies targeting highly conserved Env regions with a small CD4-mimetic achieves potent ADCC activity

**DOI:** 10.1101/2024.06.07.597978

**Authors:** Lorie Marchitto, Jonathan Richard, Jérémie Prévost, Alexandra Tauzin, Derek Yang, Ta-Jung Chiu, Hung-Ching Chen, Marco A. Díaz-Salinas, Manon Nayrac, Mehdi Benlarbi, Guillaume Beaudoin-Bussières, Sai Priya Anand, Katrina Dionne, Étienne Bélanger, Debashree Chatterjee, Halima Medjahed, Catherine Bourassa, William D. Tolbert, Beatrice H. Hahn, James B. Munro, Marzena Pazgier, Amos B. Smith, Andrés Finzi

## Abstract

The majority of naturally-elicited antibodies against the HIV-1 envelope glycoproteins (Env) are non-neutralizing (nnAbs), because they are unable to recognize the Env timer in its native “closed” conformation. Nevertheless, it has been shown that nnAbs have the potential to eliminate HIV-1-infected cells by Antibody-Dependent Cellular Cytotoxicity (ADCC) provided that Env is present on the cell surface in its “open” conformation. This is because most nnAbs recognize epitopes that become accessible only after Env interaction with CD4 and the exposure of epitopes that are normally occluded in the closed trimer. HIV-1 limits this vulnerability by downregulating CD4 from the surface of infected cells, thus preventing a premature encounter of Env with CD4. Small CD4-mimetics (CD4mc) sensitize HIV-1-infected cells to ADCC by opening the Env glycoprotein and exposing CD4-induced (CD4i) epitopes. There are two families of CD4i nnAbs, termed anti-cluster A and anti-CoRBS Abs, which are known to mediate ADCC in the presence of CD4mc. Here, we performed Fab competition experiments and found that anti-gp41 cluster I antibodies comprise a major fraction of the plasma ADCC activity in people living with HIV (PLWH). Moreover, addition of gp41 cluster I antibodies to cluster A and CoRBS antibodies greatly enhanced ADCC mediated cell killing in the presence of a potent indoline CD4mc, CJF-III-288. This cocktail outperformed broadly-neutralizing antibodies and even showed activity against HIV-1 infected monocyte-derived macrophages. Thus, combining CD4i antibodies with different specificities achieves maximal ADCC activity, which may be of utility in HIV cure strategies.

**IMPORTANCE:** The elimination of HIV-1-infected cells remains an important medical goal. While current antiretroviral therapy decreases viral loads below detection levels, it does not eliminate latently infected cells which form the viral reservoir. Here, we developed a cocktail of non-neutralizing antibodies targeting highly conserved Env regions and combined it with a potent indoline CD4mc. This combination exhibited very potent ADCC activity against HIV-1-infected primary CD4+ T cells as well as monocyte-derived macrophages, suggesting its potential utility in decreasing the size of the viral reservoir.

## INTRODUCTION

The HIV-1 envelope glycoproteins (Env) mediate viral entry into the host cell by sequentially interacting with the CD4 receptor and co-receptor CCR5 or CXCR4 (1-5). Env is synthesized as a gp160 precursor in the endoplasmic reticulum (ER) where it trimerizes and gets glycosylated (6, 7). Env is subsequently cleaved into gp120 and gp41 subunits during its transit through the Golgi apparatus (8-10). Mature HIV-1 Envs are expressed at the surface of infected cells for incorporation into viral particles. Env is a trimeric protein complex that samples different conformations; the pre-fusion “closed” State-1 Env conformation has the highest energy and is preferentially adopted by most primary isolates (11). Engagement of one Env protomer with the CD4 receptor triggers the transition from State-1 to a partially open intermediate conformation State-2, which decreases the energy barrier (12-14). Binding of two or three Env protomers to CD4 stabilizes the fully “open” State-3 CD4-bound Env conformation (14). These Env conformations are targeted by different families of monoclonal antibodies (mAbs) isolated from People Living With HIV (PLWH). Broadly-neutralizing antibodies (bNAbs) preferentially bind the “closed” State-1 conformation (11, 15-17), while non-neutralizing antibodies (nnAbs) bind normally occluded epitopes that become exposed only after Env adopts the CD4 induced (CD4i) “open” conformation (18, 19). While few individuals develop potent bNAbs, nnAbs are readily elicited upon infection and are present in most PLWH (19-23). Since nnAbs mediate potent antibody-dependent cellular cytotoxicity (ADCC) upon binding to “open” Env (19, 24-28), one strategy is to use CD4-mimetic compounds (CD4mcs) to induce conformational changes in Env similar to those that occur upon CD4 binding. These small molecule inhibitors were originally developed to sterically block Env - CD4 interaction. Here we show that CD4mcs are capable of “opening” Env and expose CD4i epitopes recognized by nnAbs naturally present in the plasma from PLWH (23-25, 29, 30), thus allowing the ADCC-mediated elimination of HIV-1-infected cells.

The opening of Env by CD4mc is a multi-step process, where the initial contact of CD4mc within the Phe43 cavity of the gp120 unmasks the co-receptor binding site (CoRBS) (24). Binding of anti-CoRBS Abs induces further conformational changes, exposing the inner domain layers 1 and 2 of gp120, which are recognized by anti-cluster A Abs (24, 31-35). The combination of the indane CD4mc BNM-III-170, anti-CoRBS Abs and anti-cluster A Abs stabilizes an asymmetric intermediate Env conformation, State2A (36), which was associated with increased ADCC responses *in vitro* and Fc-effector functions *in-vivo* (24, 36, 37). In humanized mice, this cocktail was shown to reduce the size of HIV-1 reservoir and delay viral rebound after antiretroviral therapy interruption (ARTi) (37).

An indoline-based CD4mc, CJF-III-288, with superior neutralization and ADCC activities compared to indane-based CD4mc was recently generated (38). We also identified anti-gp41 cluster I antibodies as an additional family of ADCC mediating nnAbs in the plasma from PLWH (22, 23). Here, we tested a combination of anti-cluster A, anti-CoRBS and anti-gp41 cluster I mAbs together with CJF-III-288 and found that this cocktail outperformed all previous ones with respect to eliminating HIV-1-infected primary CD4+ T cells by ADCC. Remarkably, this combination also had greater ADCC activity than a panel of well-characterized bNAbs and was able to eliminate HIV-1-infected monocyte-derived macrophages (MDMs). Detailed mechanistic analysis by smFRET imaging of Env conformations showed that this cocktail destabilized State 1 and promoted downstream open conformations, including State2A, which is known to support ADCC activity by anti-cluster A mAbs (36). The extent of conformational changes was greater than what was reported for previous nnAb cocktails, further supporting the link between the degree of “Env openness” and ADCC.

## MATERIALS AND METHODS

### Ethics statement

Written informed consent was obtained from all study participants, and the research adhered to the ethical guidelines of CRCHUM and was reviewed and approved by the CRCHUM Institutional Review Board (Ethics Committee approval number MP-02-2024-11734). The research adhered to the standards indicated by the Declaration of Helsinki. All participants were adults and provided informed written consent prior to enrollment, in accordance with the Institutional Review Board approval.

### Primary cells

CD4 T lymphocytes were purified from resting peripheral blood mononuclear cells (PBMCs) by negative selection and activated as previously described (25). Briefly, PBMCs were obtained by leukapheresis from 5 healthy non-infected adults (4 males, 1 female). CD4+ T lymphocytes were purified using immunomagnetic beads as per the manufacturer’s instructions (StemCell Technologies). CD4+ T lymphocytes were activated with phytohemagglutinin-L (PHA-L, 10 μg/ mL) for 48 hours and then maintained in RPMI 1640 complete medium supplemented with recombinant IL-2 (100 U/mL). MDMs growing was performed as previously described (39). Briefly, PBMCs were thawed, and monocytes were isolated by plate adherence in 10 cm petri dishes (Sarstedt) for 30 min in Iscove’s modified Dulbecco medium (IMDM). Non-adherent cells were collected while adherent cells were washed extensively in serum free media and allowed to differentiate to macrophages for seven days in IMDM supplemented with 100 U/mL of penicillin-streptomycin and 10% heat-inactivated pooled human sera (Valley Biomedicals), with a half media change at day 3 post-isolation.

### Plasmids and proviral constructs

Infectious molecular clones (IMCs) of the Transmitted/Founder (TF) viruses CH058, CH077, and CH040 were previously described (25, 40-43). IMC encoding HIV-1 reference strains JR-FL, JR-CSF and AD8 were described elsewhere (44-46). The vesicular stomatitis virus G (VSV-G)-encoding plasmid was previously described (47)

### Viral production, infections and *ex-vivo* amplification

For *in vitro* infection, vesicular stomatitis virus G (VSV-G)-pseudotyped HIV-1 viruses were produced by co-transfection of HEK293T cells with an HIV-1 proviral construct and a VSV-G-encoding vector using the PEI reagent (Polysciences). Two days post-transfection, cell supernatants were harvested, clarified by low-speed centrifugation (300 × g for 5 min), and concentrated by ultracentrifugation at 4°C (100,605 × g for 1 h) over a 20% sucrose cushion. Pellets were resuspended in fresh RPMI, and aliquots were stored at −80°C until use. To achieve a similar level of infection in primary CD4+ T cells among the different IMCs tested, VSV-G-pseudotyped HIV-1 viruses were produced and titrated as previously described (18, 25). Viruses were then used to infect activated primary CD4+ T cells from healthy HIV-1 negative donors by spin infection at 800 × *g* for 1 h in 96-well plates at 25°C. To expand endogenously infected CD4+ T cells, primary CD4+ T cells were isolated from PBMCs obtained from ART-treated HIV-1-infected individuals by negative selection. Purified CD4+ T cells were activated with PHA-L at 10 μg/mL for 48 h and then cultured for up to 14 days in RPMI 1640 complete medium supplemented with rIL-2 (100 U/mL) to reach greater than 5% infection for the ADCC assay. All experiments using VSV-G-pseudotyped HIV-1 isolates or *ex-*vivo amplifications were done in a biosafety level 3 laboratory following manipulation protocols accepted by the CRCHUM Biosafety Committee, which respects the requirements of the Public Health Agency of Canada.

### Antibody production

FreeStyle 293F cells (Thermo Fisher Scientific) were grown in FreeStyle 293F medium (Thermo Fisher Scientific) to a density of 1 × 10^6^ cells/mL at 37°C with 8% CO2 with regular agitation (150 rpm). Cells were transfected with plasmids expressing the light (LC) and heavy chains (HC) of a given antibody using ExpiFectamine 293 transfection reagent, as directed by the manufacturer (Thermo Fisher Scientific). One week later, the cells were pelleted and discarded. The supernatants were filtered (0.22-μm-pore-size filter), and antibodies were purified by protein A affinity columns, as directed by the manufacturer (Cytiva, Marlborough, MA, USA). Antibodies were dialyzed against phosphate-buffered saline (PBS) and stored in aliquots at −80°C. To assess purity, antibodies were loaded on SDS-PAGE polyacrylamide gels in the presence or absence of β-mercaptoethanol and stained with Coomassie blue. The anti-cluster A, anti-CoRBS and anti-gp41 Fab fragments were prepared from purified IgG (10 mg/mL) by proteolytic digestion with immobilized papain (Pierce, Rockford, IL) and purified using protein A, followed by gel filtration chromatography on a Superdex 200 16/60 column (Cytiva).

### Antibodies

The following antibodies were used to assess cell-surface Env staining and ADCC response: anti-cluster A A32 (plasmids for HC and LC were kindly provided by James Robinson); anti-CoRBS 17b (plasmids for HC and LC were kindly provided by James Robinson); anti-gp41 nnAb 246D (plasmids for HC, Cat#13741 and LC, Cat#13742 were provided by NIH AIDS Reagent Program (48)); anti-nnAb F240 (plasmids for HC and LC were previously described (49)); QA255-067 (kindly provided by Julie Overbaugh); M785U1 (50). bNAbs anti-gp41 MPER 10E8 (plasmids for HC, Cat#12290 and LC, Cat#12291 were provided by NIH AIDS Reagent Program (51)); 4E10 (plasmids for HC and LC were provided by NIH AIDS Reagent Program (52)). bNAb N6 (plasmids for HC, Cat#12967 and LC, Cat#12966 were provided by NIH AIDS Reagent Program (53)); VRC01(plasmids for HC, Cat#12035 and LC, Cat#12036 were provided by NIH AIDS Reagent Program (54)); PGT121 was provided by IAVI (55); 3BNC117 and 10-1074 (plasmids for HC and LC were kindly provided by Michel C. Nussenzweig (56, 57)). The L234A/L235A (LALA) mutations were introduced in the HC plasmids of A32, 17b, and 246D using the QuikChange II XL site-directed mutagenesis protocol. Goat anti-human IgG (H + L) (Thermo Fisher Scientific) or Goat anti-Human IgG Fc recombinant (ThermoFisher Scientific) antibodies were pre-coupled to Alexa Fluor 647 and used as secondary antibody in flow cytometry experiments. The panel of anti-HIV antibodies were conjugated with AF647 probe (Sigma Aldrich) as per the manufacturer instructions and used for cell-surface staining of HIV-1-infected MDMs. Mouse anti-human CD4 (Clone OKT4, FITC-conjugated; Biolegend, San Diego, CA, USA) and anti-p24 mAb (clone KC57; PE-Conjugated; Beckman Coulter) or Mouse anti-human CD4 (Clone OKT4, PE-conjugated; Biolegend, San Diego, CA, USA) and anti-p24 mAb (clone KC57; FITC-Conjugated; Beckman Coulter) were used to identify the productively-infected cells as previously described (58).

### Small CD4-mimetics

The small-molecule CD4-mimetic compound (CD4mc) CJF-III-288 was synthesized as described previously (38). The compound was dissolved in dimethyl sulfoxide (DMSO) at a stock concentration of 10 mM and diluted in phosphate-buffered saline (PBS) for cell-surface staining or in RPMI-1640 complete medium for ADCC assays.

### Flow cytometry analysis of cell-surface staining

Cell-surface staining of infected primary CD4+ T cells was performed 48h post-infection, as previously described (24, 25). Infected CD4+ T cells were incubated for 30 min at 37°C with anti-Env mAbs (5 µg/mL) or with plasma (dilution 1:1000). Cells were then washed once with PBS and stained with the anti-human Alexa Fluor 647-conjugated secondary antibody (2 μg/mL), AquaVivid (1:1000) and anti-CD4 FITC or PE conjugated mouse anti-CD4 Abs (1:1000) for 20 min at room temperature. After one more PBS wash, cells were fixed in a 2% PBS-formaldehyde solution. Alternatively, cells were pre-incubated with anti-cluster A, anti-CoRBS and anti-gp41 Fab fragment at 10µg/mL in the presence of CJF-III-288 before a subsequent incubation with PLWH plasma. Alexa Fluor 647 conjugated anti-human IgG Fc secondary antibodies (Invitrogen) (1:1500) were used to measure plasma binding in this context. Infected cells were then permeabilized using the Cytofix/Cytoperm Fixation/Permeabilization Kit (BD Biosciences, Mississauga, ON, Canada) and stained intracellularly using PE-conjugated mouse anti-p24 mAb or using PE of FITC-conjugated mouse anti-p24 mAb (clone KC57; Beckman Coulter, Brea, CA, USA; 1:100 dilution). Cell-surface staining of infected primary MDMs cells was performed five days post-infection, as previously described (39). Cells were washed in PBS, incubated in 10 mM EDTA for 30 min at Room Temperature, detached and transferred to 96-well V-bottom plates (Corning; Cat # 0877126). Cells were then washed twice in PBS. Prior to staining with antibodies, macrophages were incubated with 10% human sera (Valley Biomedicals) and 2% FcBlock (Miltenyi) in FACS buffer (1% BSA, 1mM EDTA in PBS) for 10 min. Following Fc blocking, macrophages were resuspended in 1% BSA and incubated with anti-Env antibodies pre-coupled to AF647 fluorophore (Sigma-Aldrich) for 30 min at RT. Cells were then washed twice with FACS buffer, fixed with 2% Paraformaldehyde (PFA). Infected cells were then permeabilized using the Cytofix/Cytoperm Fixation/Permeabilization Kit (BD Biosciences, Mississauga, ON, Canada) and stained intracellularly using FITC-conjugated mouse anti-p24 mAb (clone KC57; Beckman Coulter, Brea, CA, USA; 1:100 dilution). The percentage of productively infected cells (p24^+^) was determined by gating on the p24+/CD4-living cell population using a viability dye staining (Aqua Vivid, Thermo Fisher Scientific). Samples were acquired on Fortessa cytometer (BD Biosciences), and data analysis was performed using FlowJo v10.5.3 (Tree Star, Ashland, OR, USA).

### FACS-based ADCC assay

Measurement of ADCC using a fluorescence-activated cell sorting (FACS)-based infected cell elimination (ICE) assay was performed at 48h post-infection. Briefly, HIV-1-infected primary cells were stained with AquaVivid viability dye and cell proliferation dye eFluor670 (Thermo Fisher Scientific) and used as target cells. Cryopreserved autologous PBMC effectors cells, stained with cell proliferation dye eFluor450 (Thermo Fisher Scientific), were added at an effector: target ratio of 10:1 in 96-well V-bottom plates (Corning, Corning, NY). Target cells were treated with either DMSO or CJF-III-288 at indicated concentrations. A 1:1000 final dilution of plasma or 5 μg/mL of anti-Env mAbs was added to appropriate wells and cells were incubated for 5 min at room temperature. The plates were subsequently centrifuged for 1 min at 300 × g, and incubated at 37°C, 5% CO2 for 5 h before being fixed in a 2% PBS-formaldehyde solution. Productively-infected cells were identified by intracellular p24 and cell-surface CD4 staining as previously described (58). Samples were acquired on Fortessa cytometer (BD Biosciences) and data analysis was performed using FlowJo v10.5.3 (Tree Star). The percentage of ADCC was calculated with the following formula: [(% of p24 +CD4-cells in Targets plus Effectors) − (% of p24 +CD4-cells in Targets plus Effectors plus plasma or nnAbs)/(% of p24 +CD4-cells in Targets) × 100] by gating on infected lived target cells.

### Pseudovirions production and fluorescent labelling for smFRET Imaging

For smFRET imaging, pseudovirions were produced using HEK293T FirB cell line (59), which have high furin expression. This cell line was a kind gift from Dr. Theodore C. Pierson (Emerging Respiratory Virus section, Laboratory of Infectious Diseases, NIH, Bethesda, MD), and was cultured in DMEM (Gibco ThermoFisher Scientific, Waltham, MA, USA) supplemented with 10% (v/v) cosmic calf serum (Hyclone, Cytiva Life Sciences, Marlborough, MA, USA), 100 U/ml penicillin, 100 µg/ml streptomycin, and 1 mM glutamine (Gibco, ThermoFisher Scientific, Waltham, MA, USA) at 37°C, 5% CO_2_.

HIV-1_JR-FL_ Env pseudo-typed virions with a single gp120 domain bearing the non-natural amino acid TCO* (SiChem GmbH, Bremen, Germany) substituting the residue N135 in V1 loop and the insertion of the A1 peptide (GDSLDMLEWSLM) in V4 loop (V4-A1) were generated, purified, and fluorescently labeled as previously described (16) with minor modifications. Briefly, pNL4-3 ΔEnv ΔRT plasmid, and a 20:1 mass ratio of wild-type HIV-1_JR-FL_ gp160 plasmid to gp160 engineered to have both an amber (TAG) stop codon substituting the N135 residue in V1 loop to introduce the non-natural amino acid TCO*, and V4-A1 peptide, were co-transfected together with the plasmids NESPylRS^AF^/hU6tRNA^Pyl^ and eRF1-E55D (60) in HEK293T FirB cells (59) and in presence of 0.5 mM TCO* as previously described (61-63). Viruses were collected 48 hours post-transfection and pelleted in DPBS over a 10% sucrose cushion at 25,000 RPM for 2 hours using a SW32Ti rotor (Beckman Coulter Life Sciences, Brea, CA, USA). Virus pellet was then resuspended in labeling buffer (50 mM HEPES pH 7.0, 10 mM CaCl_2_, 10 mM MgCl_2_), and incubated overnight at room temperature with 5 μM LD650-coenzyme A (Lumidyne Technologies, New York, NY, USA), and 5 μM acyl carrier protein synthase (AcpS), which labels the A1 peptide present in Env as described (11, 16, 36). Then, the virus was incubated with 0.5 µM Cy3-tetrazine (Jena Biosciences, Jena, Germany) for 30 min at room temperature. 60 μM DSPE-PEG2000-biotin (Avanti Polar Lipids, Alabaster, AL, USA) was then added to the labelling reaction and incubated for additional 30 min at room temperature before labeled-virus purification through ultracentrifugation for 1 hour at 35,000 RPM using a rotor SW40Ti (Beckman Coulter Life Sciences, Brea, CA, USA), at 4 °C in a 6–30% OptiPrep (Sigma-Aldrich, MilliporeSigma, Burlington, MA, USA) density gradient. Labelled pseudovirions were collected, aliquoted, analyzed by western blot, and stored at -80°C until their use in imaging experiments.

### smFRET Imaging

Labelled pseudovirions previously incubated with 100 µM CJF-III-288 (38), and 50 µg/ml of each 17b and A32 monoclonal antibodies, or with the same concentrations of CJF-III-288, 17b, A32, plus mAb 246D, for 1 hour at room temperature, were immobilized on streptavidin-coated quartz slides and imaged on a custom-built wide-field prism-based TIRF microscope (61, 64). Imaging was performed in phosphate-buffered saline (PBS) pH ∼7.4, containing 1 mM trolox (Sigma-Aldrich, St. Louis, MO, USA), 1 mM cyclooctatetraene (COT; Sigma-Aldrich, St. Louis, MO, USA), 1 mM 4-nitrobenzyl alcohol (NBA; Sigma-Aldrich, St. Louis, MO, USA), 2 mM protocatechuic acid (PCA; Sigma-Aldrich, St. Louis, MO, USA), and 8 nM protocatechuate 3,4-deoxygenase (PCD; Sigma-Aldrich, St. Louis, MO, USA) to stabilize fluorescence and remove molecular oxygen. When indicated, concentrations of CJF-III-288 and mAbs were maintained during imaging. smFRET data were collected using Micromanager v2.0 (65) at 25 frames/sec. smFRET data were processed and analyzed using the SPARTAN software (https://www.scottcblanchardlab.com/software) in Matlab (Mathworks, Natick, MA, USA) (66). smFRET traces were identified according to criteria previously described (36), and traces meeting those criteria were then verified manually. Traces from each of three technical replicates were then compiled into FRET histograms and the mean probability per histogram bin ± standard error were calculated. Traces were idealized to a five-state HMM (four nonzero-FRET states and a 0-FRET state) using the maximum point likelihood (MPL) algorithm (67) implemented in SPARTAN as previously described (36). The idealizations were used to determine the occupancies (fraction of time until photobleaching) in each FRET state, and construct Gaussian distributions of each FRET state, which were overlaid on the FRET histograms to visualize the results of the HMM analysis. The distributions in occupancies were used to construct violin plots in Matlab, as well as calculate mean occupancy and standard errors. Statistical significance measures (*p*-values) of FRET state occupancies were determined by one-way ANOVA in Matlab (The MathWorks, Waltham, MA, USA). *p*-values <0.05 were considered to indicate statistical significance.

### Statistical analysis

Statistics were analyzed using GraphPad Prism version 9.1.0 (GraphPad, San Diego, CA, USA). Every data set was tested for statistical normality and this information was used to apply the appropriate (parametric or nonparametric) statistical test. P values <0.05 were considered significant; significance values are indicated as * P<0.05, ** P<0.01, *** P<0.001, **** P<0.0001.

## RESULTS

### Anti-gp41 cluster I Abs represent a major component of plasma mediated ADCC

Recent studies suggested that anti-gp41 cluster I mAbs may play a role in plasma mediated ADCC in presence of CD4mc CJF-III-288 (22, 23). To follow up on these observations, we performed Fab competition experiments. Briefly, we infected primary CD4+ T cells with HIV-1 transmitted/founder virus CH058 (CH058TF) and evaluated the capacity of plasma from 10 PLWH (Table I) to bind infected cells and mediate ADCC after pre-incubation of target cells with the CD4mc CJF-III-288 and Fab fragments of anti-CoRBS, anti-cluster A and anti-gp41 cluster I mAbs. As shown in Figure 1A, all PLWH plasma bound infected cells more efficiently upon CD4mc addition. Better binding translated into a significant improvement in ADCC (Figure 1B). In agreement with previous results (24), Fab fragments from anti-CoRBS and anti-cluster A Abs significantly decreased the capacity of PLWH plasma to mediate ADCC. However, this activity was not abrogated, suggesting the presence of additional CD4mc responsive Ab specificities in the plasma from PLWH. To test whether anti-gp41 cluster I Abs could be involved, we added a Fab fragment from F240, a well-characterized anti-gp41 cluster I nnAb recognizing the disulfide loop region (DLR) of the principal immunodominant domain (PID) of gp41 (gp120 residues 595-609) (23, 49, 68). Indeed, the combination of the three Fab fragments further decreased binding and ADCC (Figure 1A-B). These results indicated that anti-gp41 cluster I gp41Abs are responsible for a portion of plasma-mediated ADCC in PLWH.

**Figure 1.**
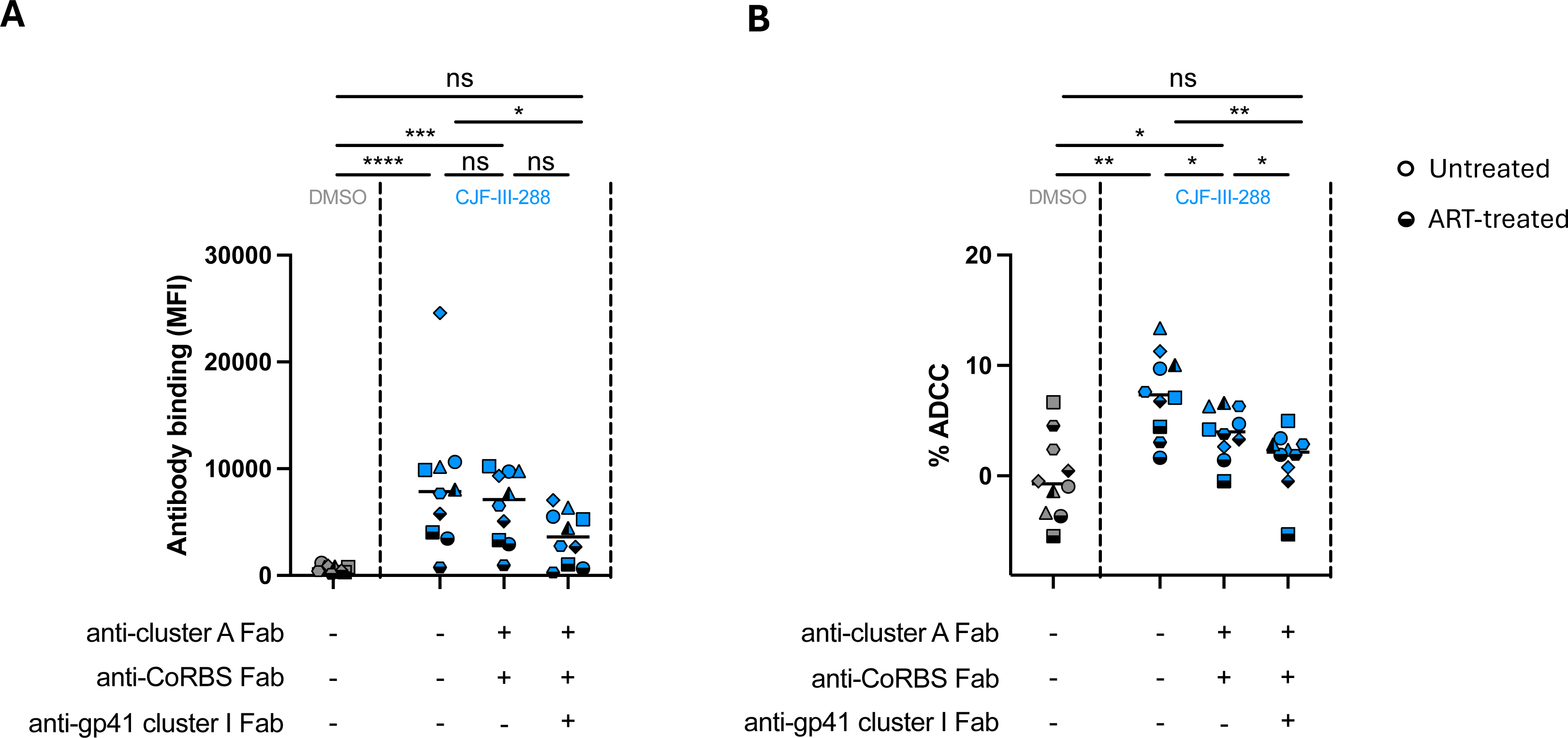
Anti-gp41 cluster I antibodies contribute to PLWH plasma-mediated ADCC. (**A**) HIV-1_CH058TF_-infected primary CD4+ T cells were pre-incubated with 10 µg/mL of each Fab antibodies in presence of CJF-III-288 depicted in blue or DMSO depicted in gray, 48h post-infection. Plasma from PLWH (dilution 1:1000) was added after incubation and plasma binding was measured by flow cytometry using Alexa-Fluor 647 conjugated anti-human IgG Fc secondary antibody. (**B**) HIV-1_CH058TF_-infected primary CD4+ T cells were used as target cells, while autologous non-infected PBMCs were used as effector cells in our FACS-based ADCC assay. Infected cells were pre-incubated with 10µg/mL of each Fab antibodies in presence of CJF-III-288 depicted in blue or DMSO depicted in gray prior to incubation with plasma from PLWH and effector cells. Each data point within a group represents an independent measurement. Median values are plotted. Statistical significance was tested using (A) Friedman test. (B) One-way ANOVA (*, P < 0.05; **, P < 0.01; ***, P < 0.001; ****, P < 0.0001; ns, non-significant).

**Table 1:**
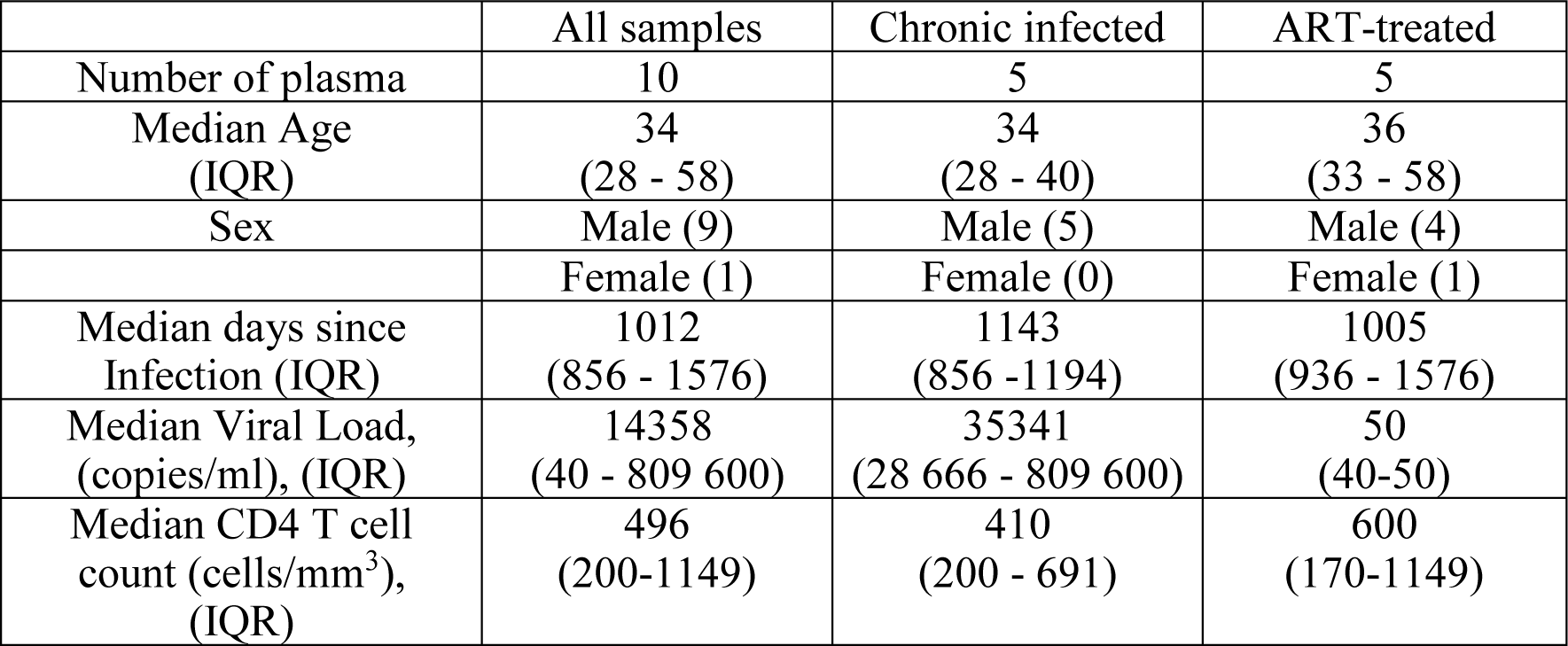
PLWH plasma, cohort characteristics, related to Figure 1.

### Development of a potent ADCC-mediating cocktail

Since the Fab blocking experiments showed that anti-CoRBS, anti-cluster A and anti-gp41 cluster I Abs all contribute to PLWH plasma-mediated ADCC in presence of CD4mc, we reasoned that a combination of monoclonal antibodies with these specificities could result in a cocktail that potently eliminates infected cells. Since the use of the anti-cluster A A32 and anti-CoRBS 17b Ab together with CD4mc was previously reported (24, 36, 37), we added anti-gp41 cluster I mAbs to the cocktail. Specifically, we tested four well-characterized anti-gp41 cluster I Abs (F240, 246D, QA255.067, M785U1) (48, 68-70) and two anti-gp41 MPER directed mAbs (10E8, 4E10) (51, 52) in combination with A32/17b for their capacity to bind and kill HIV-1-infected cells in presence of the CD4mc CJF-III-288. Compared to A32, 17b, and CJF-III-288 alone, all of the anti-gp41 cluster I mAbs tested increased binding to infected cells (Figure 2A). However, only addition of anti-gp41 cluster I 246D mAb significantly improved the ADCC activity of the cocktail (Figure 2B). As a control, we tested two monoclonal antibodies against the MPER, 10E8 and 4E10. However, neither anti-MPER mAbs increased the binding or killing of infected cells. Since 246D was the only anti-gp41 mAb significantly improving the ADCC activity of the cocktail, all subsequent experiments were performed with this mAb.

**Figure 2.**
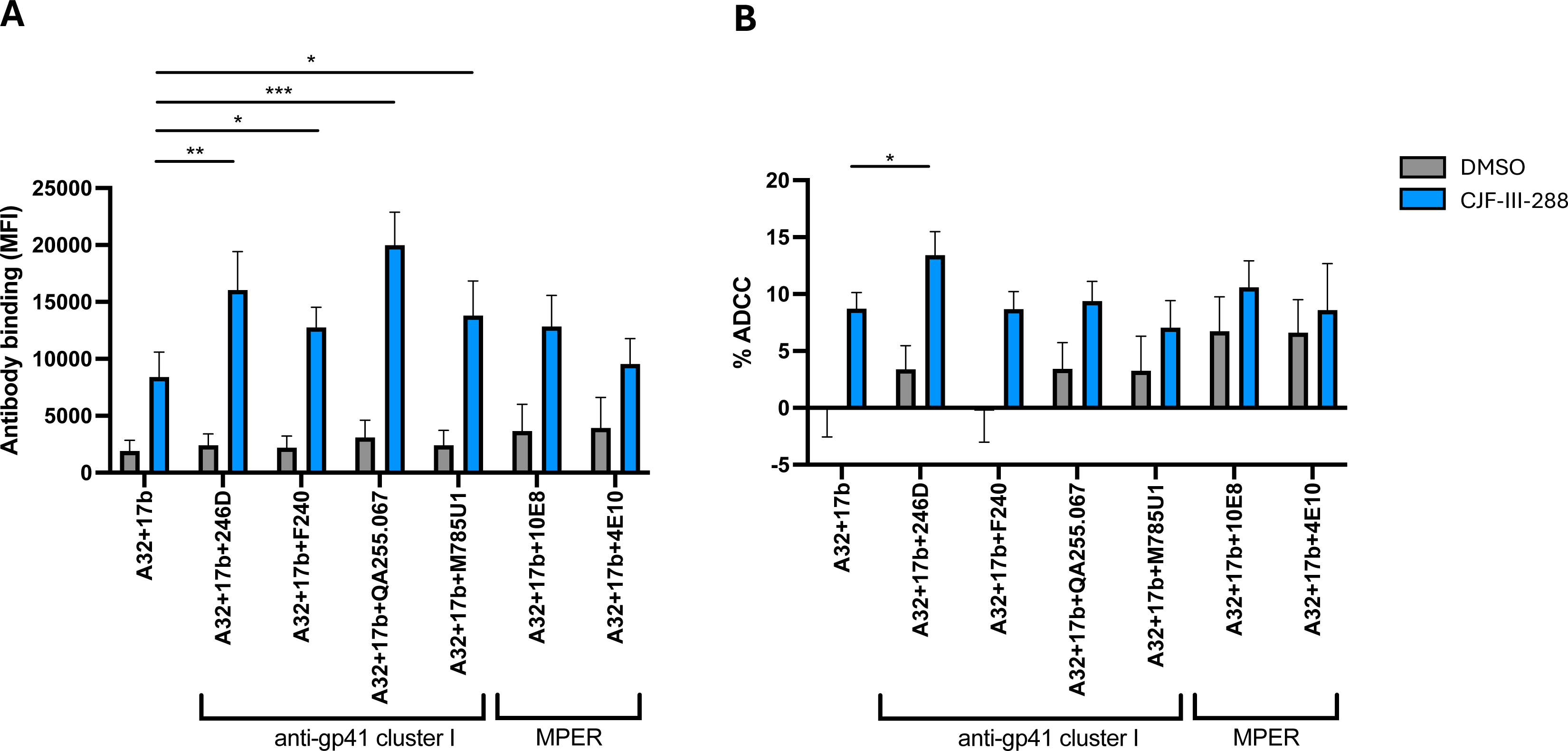
Incorporation of an anti-Cluster I mAb substantially improves the capacity of anti-cluster A/anti-CoRBS to mediate ADCC in presence of CD4mc. (**A**) HIV-1_CH058TF_-infected primary CD4 T cells were stained with a total 5µg/mL of indicated combination of nnAbs in presence of CJF-III-288 depicted in blue or DMSO depicted in gray 48h post-infection. Flow cytometry was performed to detect antibody binding using appropriated secondary antibody. The graph represents the mean fluorescence intensities (MFI) of Alexa-Fluor 647 obtained in at least 5 independent experiments. (**B**) HIV-1_CH058TF_-infected primary CD4 T cells were as target cells, while autologous non-infected PBMCs were used as effector cells in our FACS-based ADCC assay in the presence of 5µg/mL of indicated combination of nnAbs. The graph represents the percentage of ADCC obtained in presence of indicated combination of antibodies in at least 5 independent experiments. Statistical significance was tested using (A-B) Mixed-effects analysis (*, P < 0.05; **, P < 0.01; ***, P < 0.001; ****, P < 0.0001; ns, nonsignificant).

To test the combination of A32/17b/246D on a larger panel of viruses, we examined primary CD4+ T cells infected with six infectious molecular clones for their susceptibility to ADCC. In the absence of the CD4mc, no binding of infected cells or ADCC was observed. While addition of CJF-III-288 enhanced infected cell binding (Figure 3A) and ADCC (Figure 3B) by A32 and 17b, this enhancement was even more pronounced upon addition of 246D (Figure 3). The potency of the A32/17b/246D/CJF-III-288 combination prompted us to compare it to ADCC mediated by bNAbs currently used in preclinical and clinical trials (71-83). We infected primary CD4+ T cells with CH058TF and measured the capacity of the bNAbs or the cocktail to recognize infected cells and mediate ADCC. The new cocktail increased recognition of infected cells (Figure 4A) and yielded higher levels of ADCC than any one of the three CD4-binding site (CD4bs) or V3 glycan bNAbs tested (Figure 4B). We next tested *ex-vivo* expanded CD4+ T cells from ART-treated individuals (25). Briefly, primary CD4+ T cells were isolated from ART-treated PLWH (Table 2) and stimulated with PHA for 48 hours. Activated CD4+ T cells were maintained in culture with IL-2 and monitored for p24 expression by flow cytometry overtime. Staining and ADCC experiments were performed when the percentage of p24+ cells reached 5%. As for the IMC infected cells, we observed increased binding of *ex vivo* expanded cells from PLWH by the new cocktail compared to each of the bNAbs (Figure 4C). The ADCC activity of the new cocktail was also superior to all bNAbs tested (Figure 4D). These results indicated that the A32/17b/246D/CJF-III-288 cocktail was more efficient in mediating ADCC than some of the most potent bNAbs that recognize Env on the surface of CD4 T cells.

**Figure 3.**
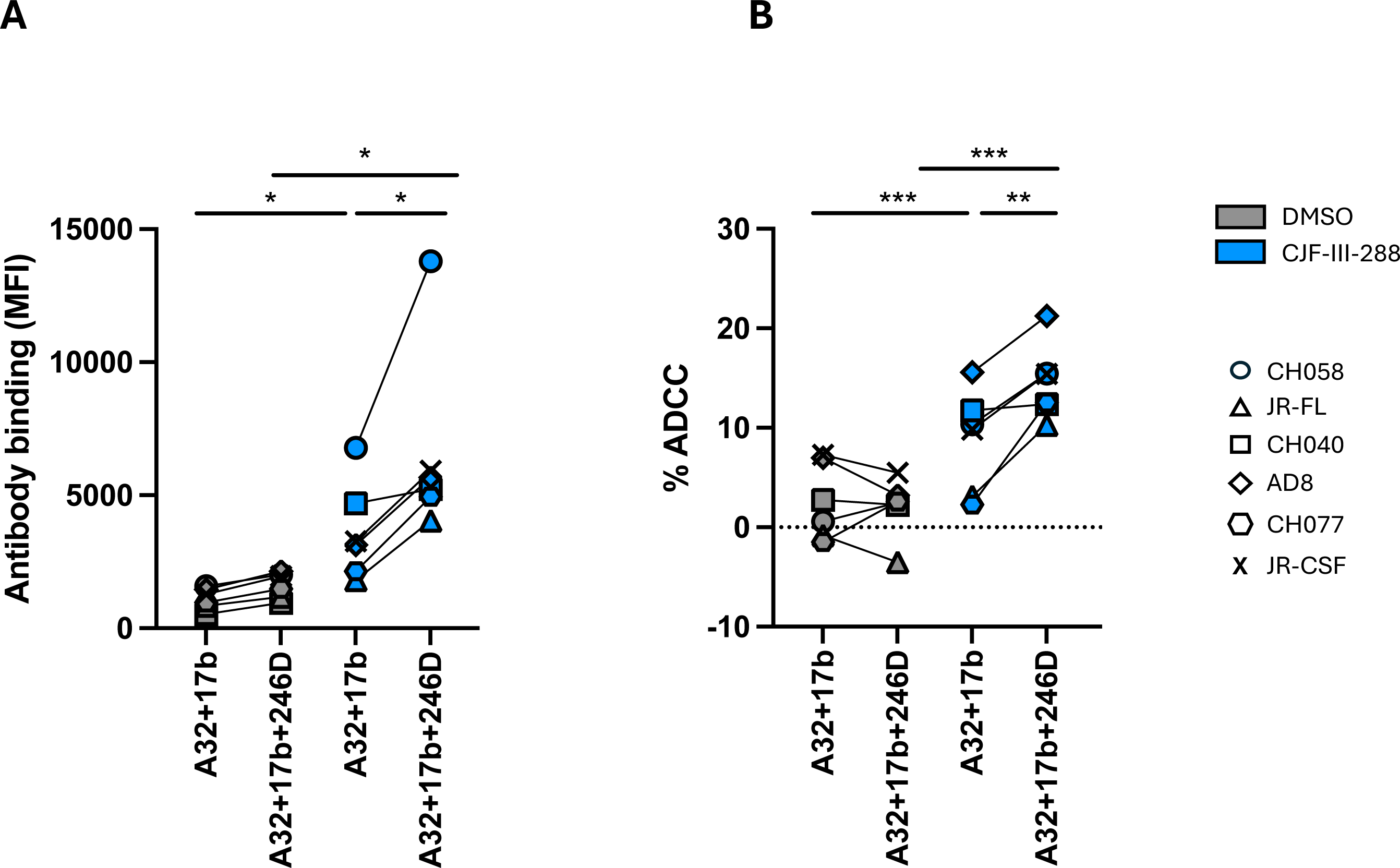
A cocktail comprising 17b, A32, 246D and CJF-III-288 mediates potent ADCC. (**A**) Primary CD4+ T cells were infected with indicated primary viruses. At 48 h post infection, cells were stained with a total 5 µg/mL of indicated antibody combination in presence of CJF-III-288 depicted in blue or DMSO depicted in gray. Flow cytometry was performed to detect antibody binding using Alexa-Fluor 647 conjugated anti-human secondary antibody. The graph represents the MeanFI of Alexa-Fluor 647 obtained in at least 3 independent experiments with each virus. Each virus is depicted as a different symbol. (**B**) Primary CD4+ T cells infected with indicated viruses were used as target cells, while autologous non-infected PBMCs were used as effector cells in our FACS-based ADCC assay in the presence of a total of 5 µg/mL of indicated combination of nnAbs. The graph represents the mean percentage of ADCC obtained from each virus in at least 3 independent experiments. Each virus is depicted as a different symbol. Statistical significance was tested using (A-B) paired t-tests or Wilcoxon tests based on statistical normality (*, P < 0.05; **, P < 0.01).

**Figure 4.**
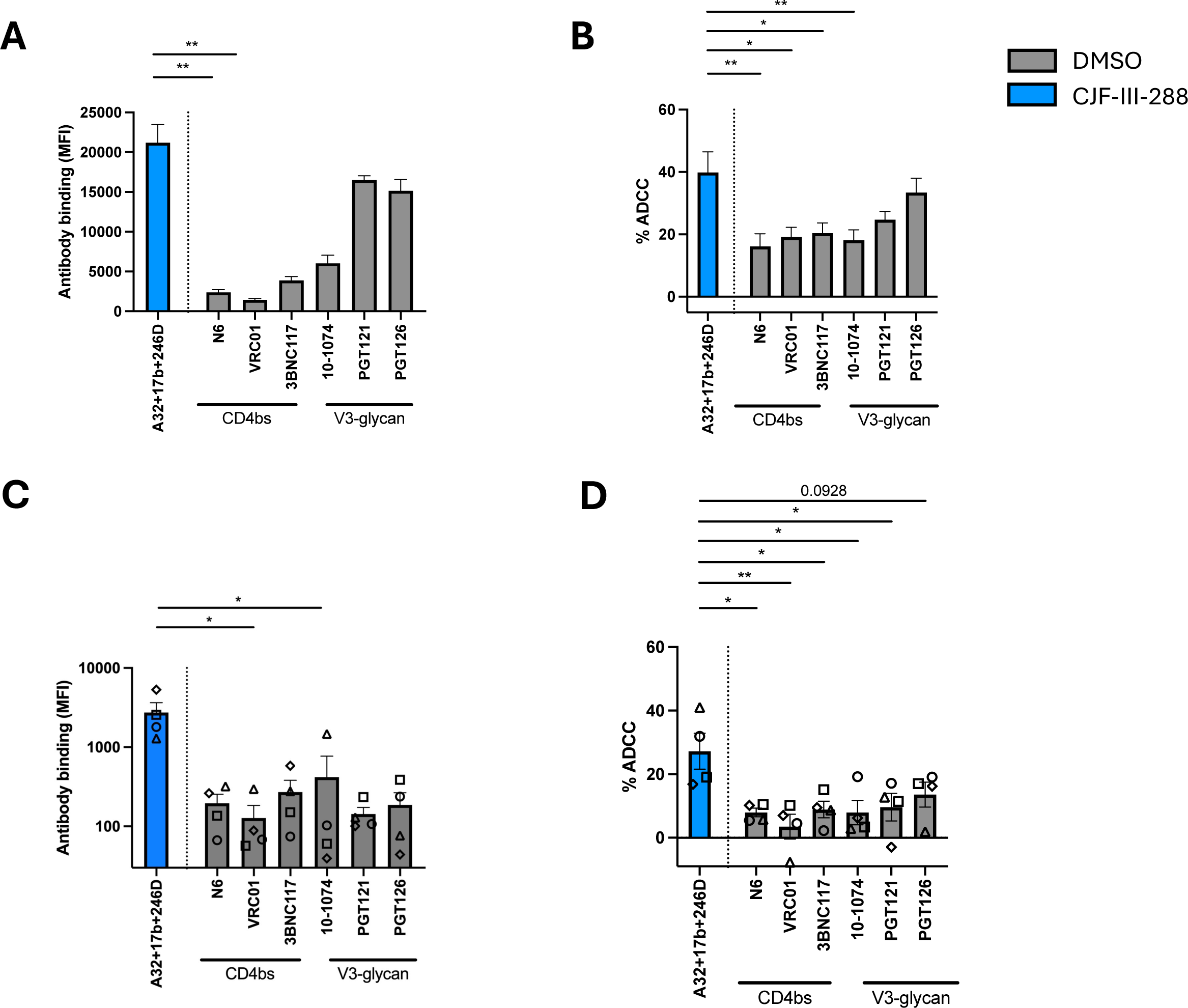
The A32/17b/246D/CJF-III-288 cocktail mediates ADCC more efficiently than bNAbs. (**A**) HIV-1_CH058TF_-infected primary CD4+ T cells were stained with 5 µg/mL of total antibodies in presence of either DMSO depicted in gray or CJF-III-288 depicted in blue 48hrs post-infection. Flow cytometry was performed to detect antibody binding. The graph represents the MFI of Alexa-Fluor 647. (**B**) HIV-1_CH058TF_-infected primary CD4+ T cells were incubated with 5 µg/mL of total antibodies in presence of either DMSO depicted in gray or CJF-III-288 depicted in blue. CD4+ T cells were used as target cells, while autologous non-infected PBMCs were used as effector cells in our FACS-based ADCC assay. The graph represents the percentage of ADCC obtained in presence of the indicated antibodies in at least 4 independent experiments. (**C**) Cell-surface staining of primary CD4+ T cells isolated from 4 HIV-1-infected individuals under ART after *ex-vivo* expansion with 5 µg/mL of total antibodies in presence of either DMSO depicted in gray or CJF-III-288 depicted in blue. Each symbol represents a different donor. Flow cytometry was performed to detect antibody binding. The graph represents the MFI of Alexa-Fluor 647. (**D**) ADCC was assessed on primary CD4+ T cells isolated from 4 HIV-1-infected individuals under ART after *ex-vivo* expansion. CD4+ T cells were used as target cells, while autologous PBMCs were used as effector cells in our FACS-based ADCC assay with 5 µg/mL of total antibodies in presence of either DMSO depicted in gray or CJF-III-288 depicted in blue. The graph represents the percentage of ADCC obtained in presence of indicated antibodies. Statistical significance was tested using (A, C) Kruskal-Wallis (B, D) One-way ANOVA, according to population normality (*, P < 0.05; **, P < 0.01).

**Table 2:**
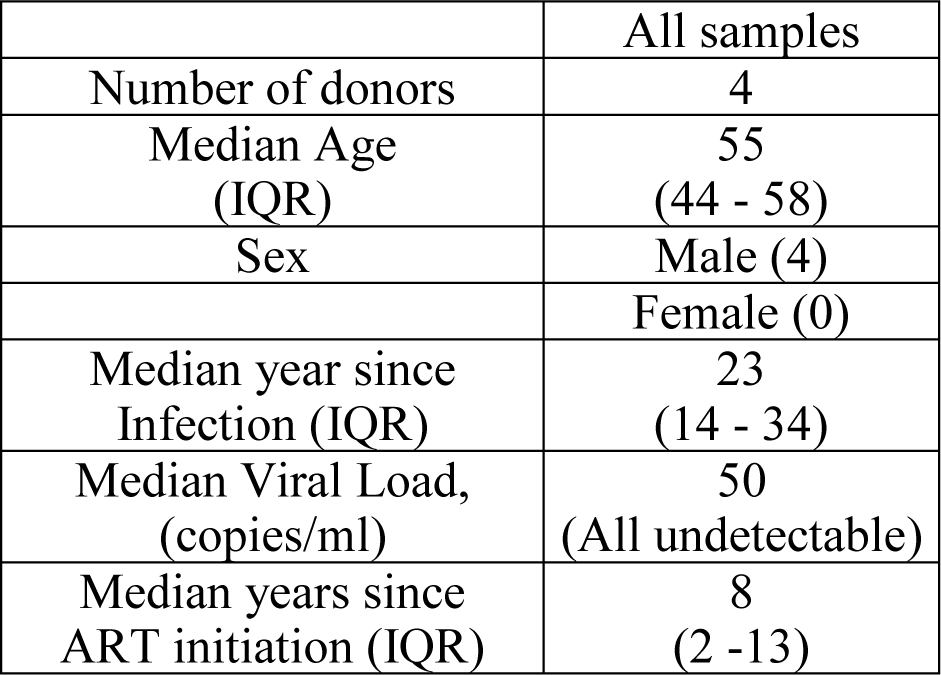
Demographic characterization of PLWH used to expand primary infected CD4 T cells, related to Figure 4.

We next assessed the relative contribution of each antibody by introducing a leucine to alanine substitution (LALA mutation) in their Fc fragment at position 234 and 235 of their heavy-chains, which is known to reduce the ability of Abs to engage Fcγ-receptors and thus abrogate ADCC (84-88). As expected, introduction of the LALA mutations did not alter the capacity of the different combination of nnAbs to bind infected cells in the presence of CD4mc (Figure 4A). In contrast, introduction of the LALA mutations into the three mAbs almost completely abrogated ADCC (Figure 5B). From the three nnAbs of the cocktail, 246D appeared to contribute to most of the ADCC activity observed. Indeed, among the various iterations tested, the biggest drop in ADCC activity was observed with the combination A32/17b/246D LALA (Figure 5B). Altogether, this data indicates that while all three nnAbs contribute to the ADCC activity of the cocktail, 246D plays a major role.

**Figure 5.**
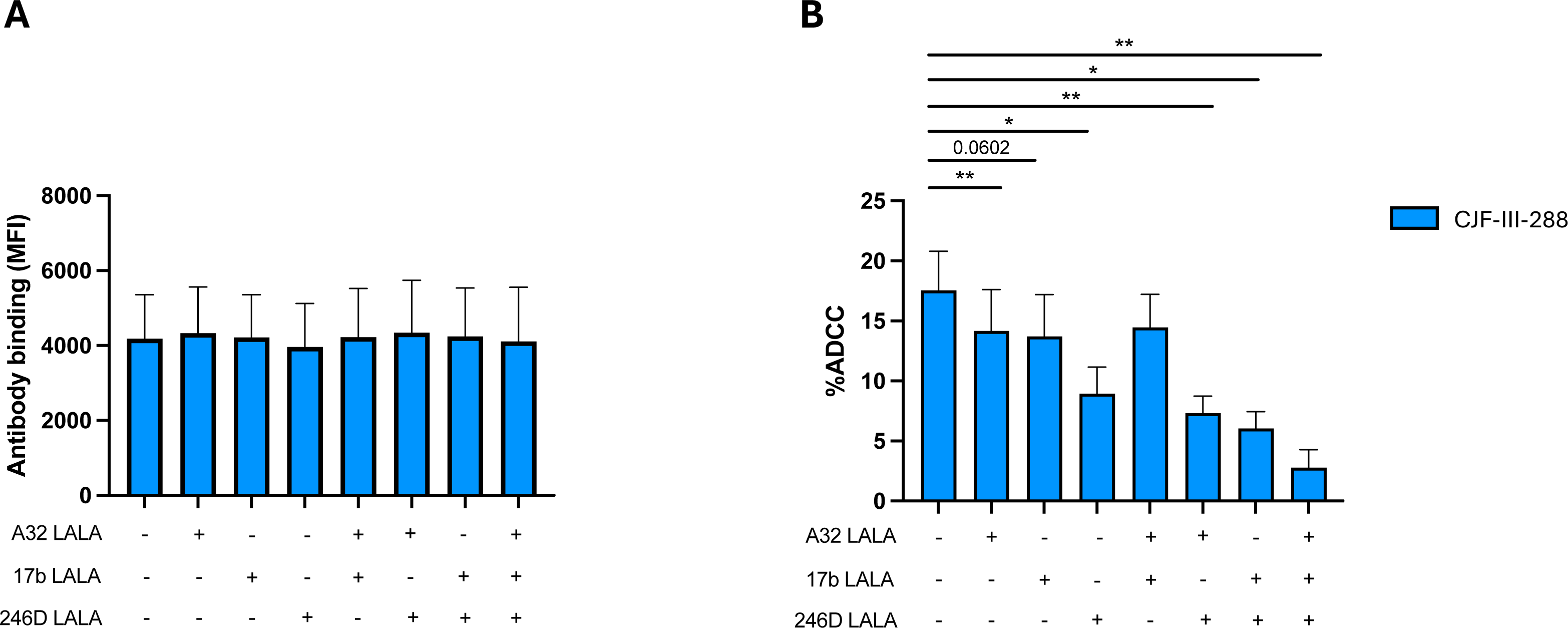
The Fc portion of all cocktail mAbs contributes to ADCC. (**A**) HIV-1_CH058TF_-infected primary CD4+ T cells were stained with a total of 5 µg/mL of indicated antibody combination in presence of CJF-III-288 depicted in blue 48h post-infection. Flow cytometry was performed to detect antibody binding using appropriated secondary antibody. The graph represents the MFI of Alexa-Fluor 647 obtained in at least 6 independent experiments. (**B**) HIV-1_CH58TF_-infected primary CD4+ T cells were used as target cells, while autologous non-infected PBMCs were used as effector cells in our FACS-based ADCC assay in the presence of 5 µg/mL of indicated combination of nnAbs. The graph represents the percentage of ADCC obtained in presence of indicated combination of antibodies in at least 6 independent experiments. Statistical significance was tested using (A-B) Wilcoxon test or paired t-test according to normality. (*, P < 0.05; **, P < 0.01).

### ADCC against HIV-1-infected monocytes-derived macrophages (MDMs)

MDMs constitute a cell lineage susceptible to HIV-1 infection *in vitro* (89, 90) and have been suggested to contribute to the viral reservoir (91-95). We previously have shown that the indane CD4mc BNM-III-170 sensitizes HIV-1-infected MDMs to ADCC mediated by plasma from PLWH (39). Given the increased potency of CJF-III-288, we evaluated the binding and ADCC activity of the new cocktail against HIV-1-infected MDMs *in vitro* (Figure 6). Briefly, MDMs were cultured for six days before infection with AD8 WT virus. Infected MDMs were stained and used as target cells 5 days post-infection. We observed that the combination of A32/17b/246D only bound to infected MDMs upon addition of the CJF-III-288 CD4mc (Figure 6B). This binding resulted in potent ADCC (Figure 6C).

**Figure 6.**
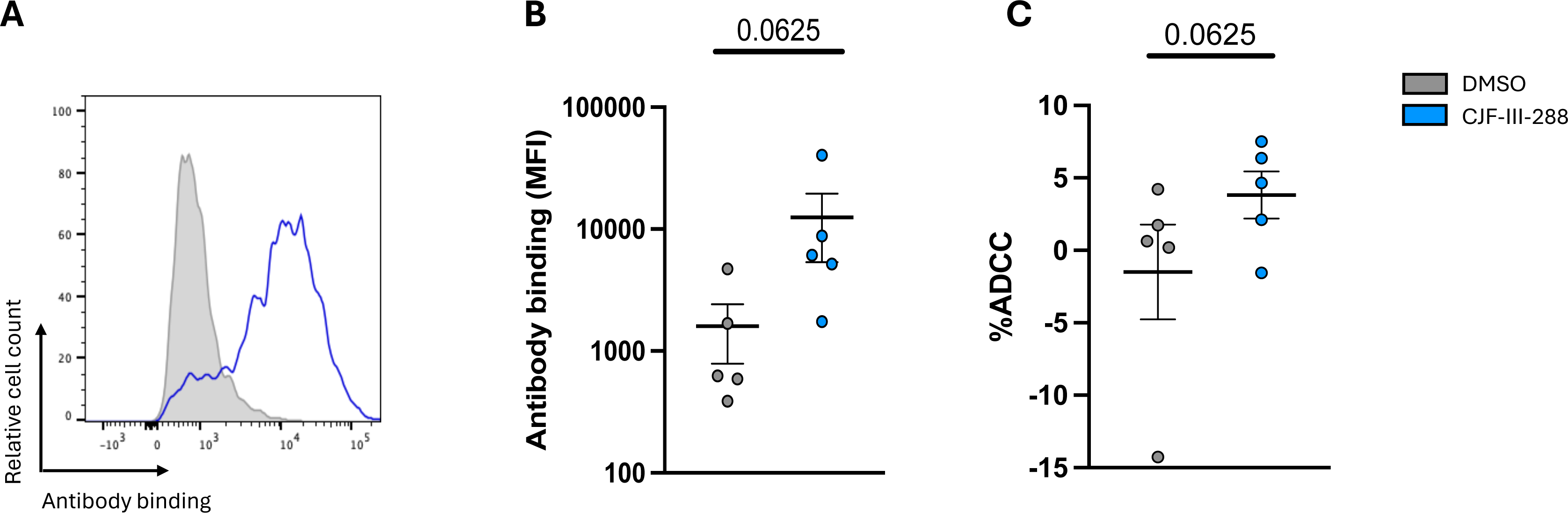
Binding and elimination of HIV-1 infected MDMs by ADCC. (**A**) HIV-1_AD8_-infected primary MDMs were stained with a total of 5 µg/mL total of the combined antibodies in presence of CJF-III-288 depicted in blue or DMSO depicted in gray 5 days post-infection. Flow cytometry was performed to detect antibody binding. (**B**) The graph represents the MFI of Alexa-Fluor 647 obtained in at least 5 independent experiments. (**C**) HIV-1_AD8_-infected primary MDMs were used as target cells, while autologous non-infected PBMCs were used as effector cells in our FACS-based ADCC assay in the presence of 5 µg/mL of indicated combination of nnAbs. The graph represents the percentage of ADCC obtained in presence of indicated combination of antibodies in at least 5 independent experiments. (B-C) Wilcoxon test or paired t-test according to normality. *p-*values are plotted.

### Impact of the combination of 17b/A32/246D/CJF-III-288 on Env conformation

We next evaluated the conformation of full-length, native Env in the presence of the 17b/A32/246D/CJF-III-288 cocktail. To this end, we applied a modified version of a well-established smFRET imaging assay (11). We attached one fluorophore to the V1 loop of gp120 using amber stop codon suppression to introduce a non-natural amino acid, followed by copper-free click chemistry with a tetrazine-conjugated fluorophore (96). The second fluorophore was enzymatically attached to the A1 peptide inserted in the V4 loop of gp120, as before (see Material and Methods) (11). Virions incorporating a single fluorescently labeled protomer among the otherwise wild-type distribution of Envs were immobilized on quartz microscope slides and imaged using total internal reflection fluorescence (TIRF) microscopy (Figure 7A). As previously reported, smFRET data indicated a predominant low-FRET conformation, consistent with the State 1. Incubation of the virions with CJF-III-288, A32, and 17b led to a dramatic destabilization of State 1 and a shift to downstream open conformations, including States 2 and 3, and the high-FRET State 2A, which was previously linked to anti-cluster A Ab binding and ADCC (Figure 7) (36). The magnitude of this effect was greater than previously observed for full-length Env in the presence of BNM-III-170, A32, and 17b (36) consistent with the greater potency of CJF-III-288 as compared to BNM-III-170 in sensitizing Env to ADCC (38). The additional presence of 246D, when combined with CJF-III-288, A32, and 17b, had a modest impact on Env conformation as shown by smFRET analysis. We observed only a slight further increase in State 2A, which did not reach statistical significance (Figure 7). This suggests that 246D does not significantly remodel gp120 conformation but may still exert an influence on gp41 conformation that is not detected with the current fluorophore attachment sites.

**Figure 7.**
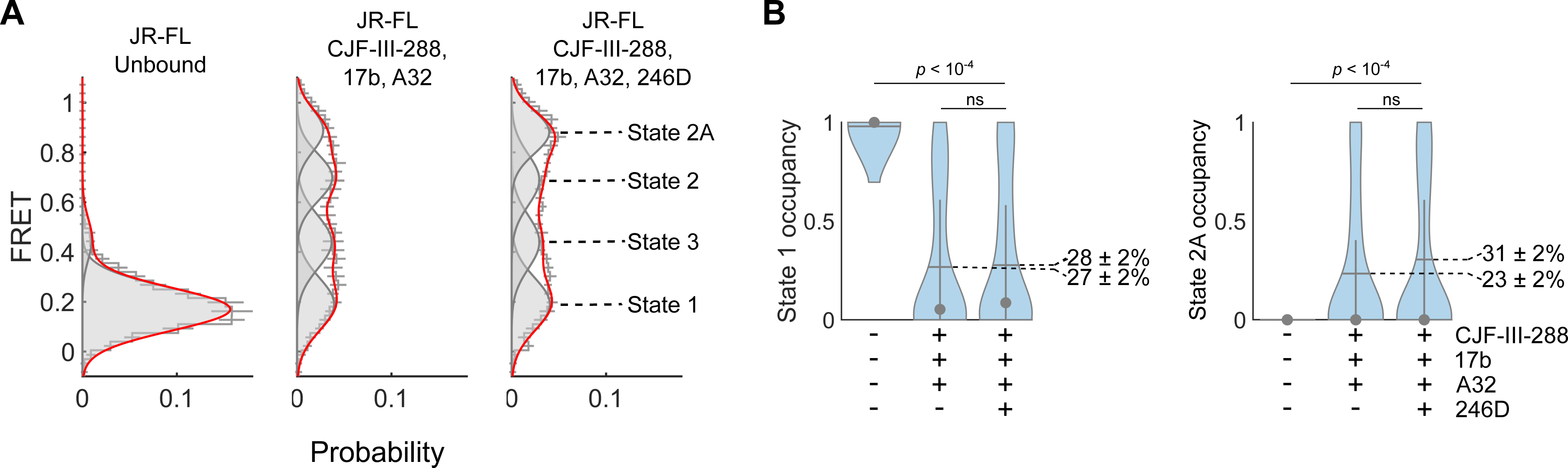
smFRET imaging of Env conformations. (A) Histograms of FRET values compiled from the population of individual Env trimers on the virion surface. smFRET data were fit using hidden Markov modeling (HMM) to a model with four non-zero FRET states. Overlaid on the FRET histograms are four Gaussian distributions (grey lines) with means and standard deviations determined through the HMM analysis. The red line indicates the sum of the four Gaussians. The histograms reflect the mean of three independent groups of trajectories with error bars corresponding to the standard error. (**B**) The occupancies in States 1 and 2A were calculated from the HMM analysis for each trace and represented with violin plots. Horizontal lines indicate the mean occupancies, while the grey circles and vertical whiskers indicate the medians and quantiles, respectively. *p*-values were determined by ANOVA.

## DISCUSSION

Despite the success of ART in suppressing viral loads, the establishment of the viral reservoir leads to a life-long infection, with increased co-morbidities. Monoclonal antibodies (mAbs) represent an attractive therapeutic approach to purge the viral reservoir due to their capacity to eliminate infected cells by Fc-effector functions, including ADCC. One strategy is to harness the ADCC activity of nnAbs by enabling their recognition of infected cells using CD4mc to “open-up” Env (24, 25, 36, 38, 39, 84, 97, 98). Here we improved a cocktail of nnAbs and CD4mc, which showed promise at reducing the size of the viral reservoir in hu-mice (37), by adding a new nnAb targeting a conserved epitope in the gp41 (23) and replacing the indane BNM-III-170 by the newly developed indoline CJF-III-288 CD4mc (38).

Addition of 246D to the previous A32/17b combination enhanced the capacity of the cocktail to eliminate HIV-1-infected cells in the presence of CD4mc. The new cocktail was also superior to the potent bNAbs N6, VRC01, 3BNC117, 10-1074, PGT121 and PGT126 at eliminating infected cells in both *in vitro* and *ex vivo* settings. Mechanistic studies by smFRET indicate that this cocktail decreases State-1 occupancy and stabilizes Env in downstream conformations which are vulnerable to ADCC (18, 19, 24-26, 28, 36, 37), although addition of 246D did not increase more open conformations compared to A32/17b. This suggest that the improved ADCC activity is due to the addition of an extra Fc-portion, from 246D, which likely helps in crosslinking the FcγR in effector cells. Indeed, while the Fc portion of all three nnAbs were found to be required to mediate potent ADCC, the anti-gp41 cluster I 246D Ab played a predominant role. Whether the Fc-portion of an Ab targeting gp41 facilitates the clustering of FcγR on effector cells remains to be demonstrated. Nonetheless, the A32/17b/246D/CJF-III-288 combination was efficient at eliminating infected MDMs, a cell type which was previously reported to be resistant to mAb-mediated ADCC (99).

The A32/17b/246D/CJF-III-288 cocktail may have therapeutic utility as it targets four independent epitopes that represent some of the most conserved regions of HIV-1 Env. Figure 8 projects the A32, 17b, 246D epitopes and CJF-III-288 binding site onto an untriggered closed Env trimer and shows the conservation of Env residues contributing to the binding of each component based on available structural information of their antigen complexes or peptide mapping. A32 and 246D epitopes are localized within the interior of the Env trimer (Figure 8A). The A32 epitope maps onto the gp120 inner domain proximal to the N- and C-termini of gp120 and around the α0- and α1-helices (31, 34, 100), while 246D binds to a linear peptide in the immunodominant cluster I region of gp41 (residues 596-606) (48). Analysis of sequence conservation among HIV-1 isolates indicates that both epitopes map to highly conserved Env regions that contain gp41-gp120 interprotomer contacts (Figure 8b). The A32 epitope in particular is located in the cluster A gp120 region) (50) and is directly involved in CD4-triggered conformational changes in the gp120 inner domain and thus constitute some of the most conserved regions of Env with certain residues or motifs being identical among divergent HIV isolates (e.g. TLFC^54^, W^69^, THACVPTDP^79^ and Q^103^, D^107^ S^110^, Y^217^ and PA^221^) (31, 34, 100). In contrast to A32 and 246D, the epitopes for 17b and the CJF-III-288 binding site map to the outer domain of gp120; 17b maps to the outer domain of gp120, proximal to the CD4 binding site (101-103) and CJF-III-288 binds within the Phe43 cavity in the CD4 binding site (38). The 17b epitope is well conserved among HIV-1 isolates with more than 65% of Env residues that form the epitope being highly conserved (101). Similarly, CJF-III-288 which targets the CD4 binding cavity interacts with strictly conserved residues of gp120 (e.g. ST^257^ FN^377^, F^382^, Y^384^, W^Q428^, and GG^473^). Based on the high sequence conservation of these epitopes among the HIV isolates, it is likely that treatments with a mix of A32/17b/246D/CJF-III-288 may be highly cross-reactive and undergo limited immune escape.

**Figure 8.**
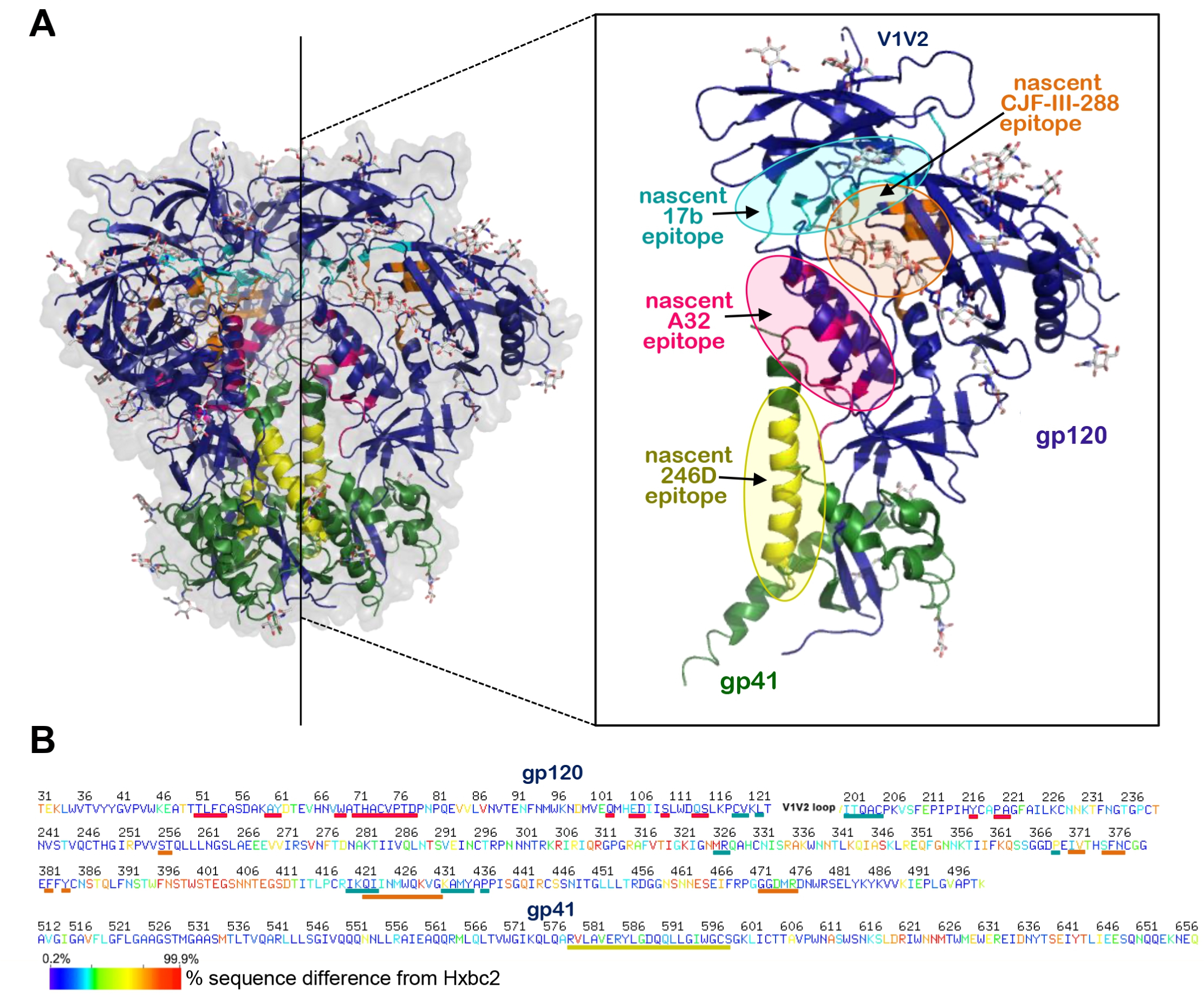
Env regions targeted by the A32/17b/246D/CJF-III-288 cocktail. (B) Location of A32 (anti-cluster A), 17b (anti-CoRBS), 246D (anti-gp41 cluster I) and CJF-III-288 (CD4mc) binding sites within the unliganded HIV-1 trimer. The BG505 SOSIP.664 gp140 unliganded trimer (PDB code: 4ZMJ) is shown, colored green and dark blue for gp41 and gp120 protomers. The nascent epitopes for A32, 17b, 246D and CJF-III-288 are colored magenta, cyan, yellow and orange, respectively. The nascent epitopes are defined as gp120 contact residues contributing buried surface area (BSA) to the Env antigen-Fab interface calculated from available structures: A32 (PDB code: 4YC2), 17b (PDB code: 6CM3) and CJF-III-288 (PDB code:8FM3). Epitope for 246D is as defined by peptide scanning in (48). The blow-up view shows insight into one gp41-gp120 protomer with epitopes. (**B**) Sequence conservation of A32, 17b, 246D and CJF-III-288 binding regions among all listed Env sequences from the Los Alamos Data Base. The sequence of gp120 and gp41, excluding the V1V2 region, is shown based on the conservation of each residue e.g. residues differed 0.2-7% and 87-99.9% from the Hxbc2 sequence colored with dark blue and red. Lines (A32: magenta, 17b: cyan, 246D: yellow and CJF-III-288: orange) below the sequence indicate the epitope footprints.

Overall, here we show that upon CD4mc addition, antibodies targeting the immunodominant cluster I region of gp41 comprise a major fraction of PLWH plasma ADCC activity. Supplementing anti-gp41 cluster I to cluster A and CoRBS antibodies greatly enhanced ADCC mediated cell killing in the presence of a potent indoline CD4mc, CJF-III-288. By combining nnAbs targeting multiple conserved epitopes we achieved broad and potent ADCC activity, which may be of utility in HIV cure strategies.

## ACKNOWLEDGMENTS

The authors thank the CRCHUM BSL3 and Flow Cytometry Platforms for technical assistance, Mario Legault from the FRQS AIDS and Infectious Diseases network for providing the human PBMCs. We thank Dennis Burton (The Scripps Research Institute) for kindly providing the infectious molecular clone JR-FL. We thank the following collaborators for kindly providing Abs: Julie Overbaugh (Fred Hutchinson Cancer Research Center) for QA255-067. The following reagents were obtained by the NIH AIDS reagents program: 246D (contributed by Dr. Susan Zolla-Pazner); 10E8 (contributed by Dr. Mark Connors); 4E10 (contributed by DAISS/NIAID); N6 (contributed by Dr. Jinghe Huang et Dr. Mark Connors); VRC01 (contributed by John Mascola). We thank Michel C. Nussenzweig for 3BNC117 and 10-1074 and James Robinson for A32 and 17b. This study was supported by a CIHR Team grant #422148, a Canada Foundation for Innovation (CFI) grant #41027 to A.F and by the National Institutes of Health to A.F. (R01 AI148379, R01 AI150322, R01AI176531), A.F. and M.P. (AI174908) and M.P. and Georgia Tomaras (P01 P01AI162242). B.H.H. is supported by R01 AI162646, UM1AI144371 and UM1AI164570. Support for this work was also provided by UM1AI164562 (ERASE) to A.F., and A.B.S. A.F. was supported by a Canada Research Chair on Retroviral Entry RCHS0235 950– 232424. J.P., and G.B.B. were supported by CIHR doctoral fellowships. M.N. was supported by a ViiV fellowship. M.B. was supported by a FRQS doctoral fellowship. A.T. was supported by a MITACs Elevation post-doctoral fellowship. The funders had no role in study design, data collection and analysis, decision to publish, or preparation of the manuscript.

## AUTHOR CONTRIBUTIONS

L.M., J.R., J.P., and A.F. conceived the study. L.M., J.R., and A.F. designed experimental approaches. L.M., J.R., J.P., A.T., M.A.DS., M.N., M.B., G.B.B., S.P.A., K.D., E.B., D.C., H.M., C.B., B.H.H., J.M. M.P. and A.F. performed, analyzed, and interpreted the experiments. D.Y., TJ.C., HC.C., B.H.H., M.P. and A.B.S., supplied novel/unique reagents. L.M., J.R., B.H.H and A.F. wrote the paper. All authors have read, edited, and approved the final manuscript.

## DISCLAIMER

The views expressed in this manuscript are those of the authors and do not reflect the official policy or position of the Uniformed Services University, US Army, the Department of Defense, or the US Government.

## CONFLICT OF INTEREST

The authors declare no competing interests.

## DATA AVAILABILITY

Data and reagents are available upon request.

## Notes

### Competing Interest Statement

The authors have declared no competing interest.

